# Balancing sample sizes in parasitology: A novel experimental infection method using faecal parasite eggs and aquatic intermediate hosts

**DOI:** 10.1101/2025.10.24.683585

**Authors:** Sara M. Rodríguez, Katherine Burgos-Andrade, Valentina Escares-Aguilera, Bárbara Gutiérrez, Nelson Valdivia

## Abstract

Balancing sample sizes between infected and uninfected hosts is a common challenge in ecological parasitology, particularly when dealing with intermediate hosts involved in complex life cycles. In these life cycles, common in nature, the intermediate hosts accumulate parasites across their ontogeny, making it difficult to obtain similar numbers of naturally infected and uninfected individuals across full host body-size spectra. Here, we present a standardized and repeatable method to experimentally infect and re-infect intermediate hosts using parasite eggs derived directly from the fresh faeces of definitive hosts. We tested this methodology in a marine host–parasite system involving the acanthocephalan *Profilicollis altmani*, the mole crab *Emerita analoga*, and the grey gull *Leucophaeus modestus*. Crabs were raised in controlled laboratory conditions from early juvenile stages, ensuring no prior infections, and exposed to parasite eggs through filtered faecal suspensions. Larval development was tracked across two inoculation events. The first larval stage (acanthellae) appeared six days after inoculation and matured into cystacanths within the following week. After the first exposure, 55% of inoculated hosts were infected; after the second, infection prevalence reached 100%. Larger crabs acquired more parasites than smaller ones, likely due to higher filtration rates. The method closely mimics natural faecal-oral transmission while avoiding the need to isolate or handle parasite eggs directly. It also overcomes the common issue of parasite overdispersion in natural populations by enabling controlled infections across host size classes and experimental replicates. This technique is particularly useful in species with opaque cuticles, where internal parasites cannot be visually identified in vivo. Additionally, it can be extended beyond acanthocephalans to other helminth taxa (e.g., nematodes, cestodes, and digeneans) whose eggs are also shed in vertebrate faeces. Our results establish a reliable, ecologically relevant protocol for experimental parasitology in marine and freshwater systems.

## INTRODUCTION

Parasites are ubiquitous in animal populations and ecosystems, often exerting strong influence on host dynamics (May & Anderson, 1978; Thomas, Guégan & Renaud, 2005). A fundamental pattern in parasite ecology is that parasites are heterogeneously distributed among hosts; that is, not all hosts have the same probability of becoming infected, nor do they carry the same parasite burden (Anderson & May, 1978; May & Anderson, 1978). This heterogeneity is even more pronounced in parasites with complex life cycles, where smaller hosts (assuming that body size correlates with host age; Ebert, 1999) are typically less parasitized than larger ones (Rodríguez & Valdivia, 2017; Rodríguez et al., 2022). This heterogeneous distribution poses challenges for sampling and replication under natural field conditions (Cattadori, Boag & Hudson, 2008; Filazzola & Cahill Jr, 2021). Conducting experiments in the field reduces the likelihood of obtaining replicable samples, in contrast to other biological disciplines that work under controlled conditions. Moreover, because hosts accumulate parasites throughout their ontogeny, collecting both parasitized and non-parasitized individuals across a full range of host sizes becomes difficult. Consequently, generating *a priori* predictions of infected hosts, rather than *a posteriori* interpretations, is nearly impossible under these conditions. In this context, manipulative experiments become essential, as they allow researchers to isolate and control the main factors of interest, such as host size and parasite infection, thereby enabling robust and sound hypothesis testing. Experimental infections thus provide a powerful framework to control host variability, stratify and balance sample sizes, and improve the statistical reliability of studies in complex host–parasite systems.

Researchers have developed various techniques to sample and analyze parasitized individuals to assess the effects of parasitism under controlled experimental conditions (Franceschi et al., 2008; Dianne et al., 2011, 2012, 2014; Thünken et al., 2010, 2018). One of the most common approaches involves extensive fieldwork to collect individuals across different ontogenetic stages, locations, and environmental conditions (Altman & Byers, 2014; Rodríguez & Valdivia, 2017; Goedknegt et al., 2019; Rodríguez et al., 2022). These individuals are then dissected to determine their infection status and parasite load (Altman & Byers, 2014; Rodríguez & Valdivia, 2017; Balboa-Figueroa et al., 2019; Rodríguez et al., 2017a, 2022). Other technique involves collecting infected and uninfected individuals from historically parasite-free areas, using them as control groups for experimental treatments (Byers et al., 2008; Goedknegt et al., 2019; Díaz-Morales et al., 2022). However, these methods present two major limitations. First, because the experiments are typically conducted without prior knowledge of individual infection status, they do not allow for *a priori* hypothesis testing. Second, it is difficult to balance the number of infected and uninfected individuals, which limits statistical power and replication. A different approach involves visually detecting parasite larvae within transparent-bodied intermediate hosts (Duclos et al., 2006; García-Varela et al., 2013; Thünken et al., 2018). For example, in species with translucent cuticles, such as the amphipod *Gammarus pulex*, it is possible to detect parasite cystacanths (i.e., mature, infective larval stage) through direct observation (Bakker et al., 1997; Thünken et al., 2010, 2018). These visual techniques help homogenize sample groups and improve replication by allowing pre-selection based on infection status. However, such approaches are only feasible in hosts with transparent exoskeletons. In most invertebrates with opaque cuticles, it is not possible to determine infection visually prior to dissection (Balboa-Figueroa et al., 2019; Rodríguez et al., 2017a, 2017b, 2022). Furthermore, in species where parasites reside within internal body cavities such as the coelom, there are currently no effective antiparasitic treatments or non-lethal methods to clear infections before experiments. In other systems, intermediate hosts have been artificially infected by directly inoculating parasites under the cuticle or carapace (Pulgar et al., 1995). While effective in inducing infection, these methods bypass the natural transmission route and may not accurately reflect ecological conditions. Therefore, developing methodologies that allow for controlled, ecologically relevant infection remains a key challenge in experimental parasitology.

Existing methodologies manipulate parasite prevalence and burden experimentally in intermediate hosts, often aiming to balance sample sizes across treatments (Mouritsen, 2002; Koprivnikar & Poulin, 2009; Larsen & Mouritsen, 2014; Díaz-Morales et al., 2022). Dianne et al. (2012) extracted parasite eggs from mature females in definitive hosts’ intestines and deposited them on intermediate hosts’ food, increasing infection probability compared to placing eggs directly in the habitat (water). However, this requires separating nearly 100 eggs per host to ensure acceptable infection success (Dianne et al., 2011, 2014). Another method involves screening hosts for parasites and inducing parasite emergence, mainly cercariae, by manipulating factors like light, temperature, and salinity (Mouritsen, 2002; Poulin, 2006; Larsen & Mouritsen, 2014). Subsequently, infected and potentially uninfected hosts are kept separately and re-infected before experiments (Koprivnikar & Poulin, 2002; Díaz-Morales et al., 2022). Nevertheless, these techniques cannot guarantee that initial deworming was successful or that reinfection results solely from experimental inoculations rather than natural infection. In parasites with complex life cycles, definitive hosts release infective stages through faeces, which intermediate hosts consume. Therefore, inoculations using faeces containing infective stages could enable better sample balancing and replication.

The use of faeces to diagnose enteroparasite infections is continually used in human, veterinary, and wildlife diseases (Mas-Coma et al., 2014; Wolf et al., 2014; De Nys et al., 2015; Rodríguez & George-Nascimento, 2021). Faecal samples are an important source of information on parasites, which is mainly used as detection technique. To our knowledge, the use of faeces as infection methodology has only been and scarcely used as treatment of faecal microbiota transplantation in humans (Messias et al., 2018; Lin et al., 2020). Even though the faecal samples have been used for treatment of infectious diseases and seem to be a good methodology for diagnosis and therapies (Presswell & Lagrue, 2016; Lin et al., 2020; Rodríguez & George-Nascimento, 2021), faecal samples have been null or scarcely used as infection technique in ecological studies (Aznar et al., 2012; Rodríguez & George-Nascimento, 2021). This scenario occurs despite of the abundant empirical research in infections dynamics, especially in parasites with complex life cycle, which have been based on *a posteriori* hypothesis formulation. In parasite with complex life cycles, definitive hosts release the infective stage through the faeces into the environment, and these are consumed by the intermediate hosts. Therefore, inoculations or experimental infections using faeces with infective stages offer the possibility to generates *a priori* hypotheses, to balance sample size, and obtain sufficient samples for appropriate replication.

Here, we propose a novel, reliable experimental methodology to inoculate faeces containing parasite eggs to intermediate hosts. We apply this methodology in a complex parasite system that includes the acanthocephalan *Profilicollis altmani*, crustaceans, the mole crab *Emerita analoga* of different body sizes, and the faeces of one of its definitive seagull hosts, the grey gull *Leucophaeus modestus*. This study had three main aims: (i) to infect intermediate hosts of different ontogenetic stages, under controlled conditions using faecal samples with parasite eggs, (ii) to generate a post-inoculation increase of parasite burden in hosts regardless of body size, and (iii) to demonstrate the capacity to balance infected and uninfected hosts across different body sizes, enabling *a priori* hypotheses about infection dynamics, balancing sample sizes, and increasing replicability under controlled conditions compared to natural field variability.

## MATERIALS AND METHODS

### 2.1 Host collection and acclimatization

Mole crabs (*Emerita analoga*) were collected from August 2018 to October 2019 in Curiñanco, southern-central Chile (39.4° S – 73.2° W). Within this temporal range, six sampling times were conducted. The mole crabs collected in the first, second, and third sampling times (August 2018, December 2018, and March 2019, respectively) were grouped into Group 1. The mole crabs collected in the fourth, fifth, and sixth sampling times (June 2019, September 2019, and November 2019) were grouped into Group 2. These groups were raised in the laboratory for different periods to obtain two groups of body sizes (Group 1 was raised for 9–12 months: final cephalothorax length 14–31.6 mm [large crabs]; Group 2 for 3–10 weeks: final size 8–13.9 mm [small crabs]). Across both groups, median and maximum cephalothorax length were 13.9 mm and 31.6 mm, respectively. The objective of using a range of dates for collecting the mole crabs in each group was to capture individuals at the beginning and end of the recruitment period, thereby obtaining specimens of varying sizes but within a roughly similar size range (Lastra et al., 2004).

In each sampling time, during low tide and with the help of plastic corers (0.03 m^-2^), which were buried to a depth of 30 cm approximately, individuals of *E. analoga* with body size less than 9 mm were captured, as the mole crabs are not parasitized by *P. altmani* below this size (Rodríguez & Valdivia, 2017). We captured 350 individuals each time, which were transported alive to laboratory of Coastal Ecology of Universidad Austral de Chile. When transported in a less-than-one-hour drive, the mole crabs were maintained in buckets filled with natural seawater, with continuous aeration supplied by a battery-powered air pump to minimize stress on the individuals.

In the laboratory, crabs were kept in 60 x 30 x 25 cm aquaria. The bottom (ca. 1/3 of aquarium height) of each aquarium consisted of sand previously cleaned with freshwater to eliminate any parasite’s living egg. Each aquarium was filled with filtered seawater (1 μm and 5 μm filters, plus a carbon and UV-C (200-280 nm) filter), and kept with constant aeration and at 15 °C. The mole crabs were fed daily with 150 ml of *Isochrysis galbana* microalgae, which were grown in Microalgae Laboratory of Universidad Austral de Chile, following the protocol of Guillard (1975). The water in each aquarium was replaced once a week. In this way, the crabs grew under cultivated conditions without external contaminant agents and neither the possibility of naturally parasitizing. Large mole crabs were acclimatized and maintained in these aquaria under bubbling seawater and ad libitum feeding conditions for 9 to 12 months. Similarly, small mole crabs were acclimatized under the same conditions, with circulating seawater and ad libitum feeding, but for a shorter period of 3 to 10 weeks. Prior to infection experiments, each size group (Large and small mole crabs) was divided into four aquaria: two were inoculated with acanthocephalan parasite eggs and microalgae, and two served as controls (microalgae only).

### 2.2 Faeces collection and eggs detection

During December 2019, at Curiñanco Beach, 15 fresh faecal samples (approximately 3–5 ml) from the grey gull *Leucophaeus modestus* were collected (Fig. 1A). Using a telescope, individuals actively foraging along the shoreline were kept in view, that is, visually followed until defecation occurred. Faeces were stored in 50 ml sterilized Falcon tubes and transported to Coastal Ecology Laboratory.

**Figure 1:**
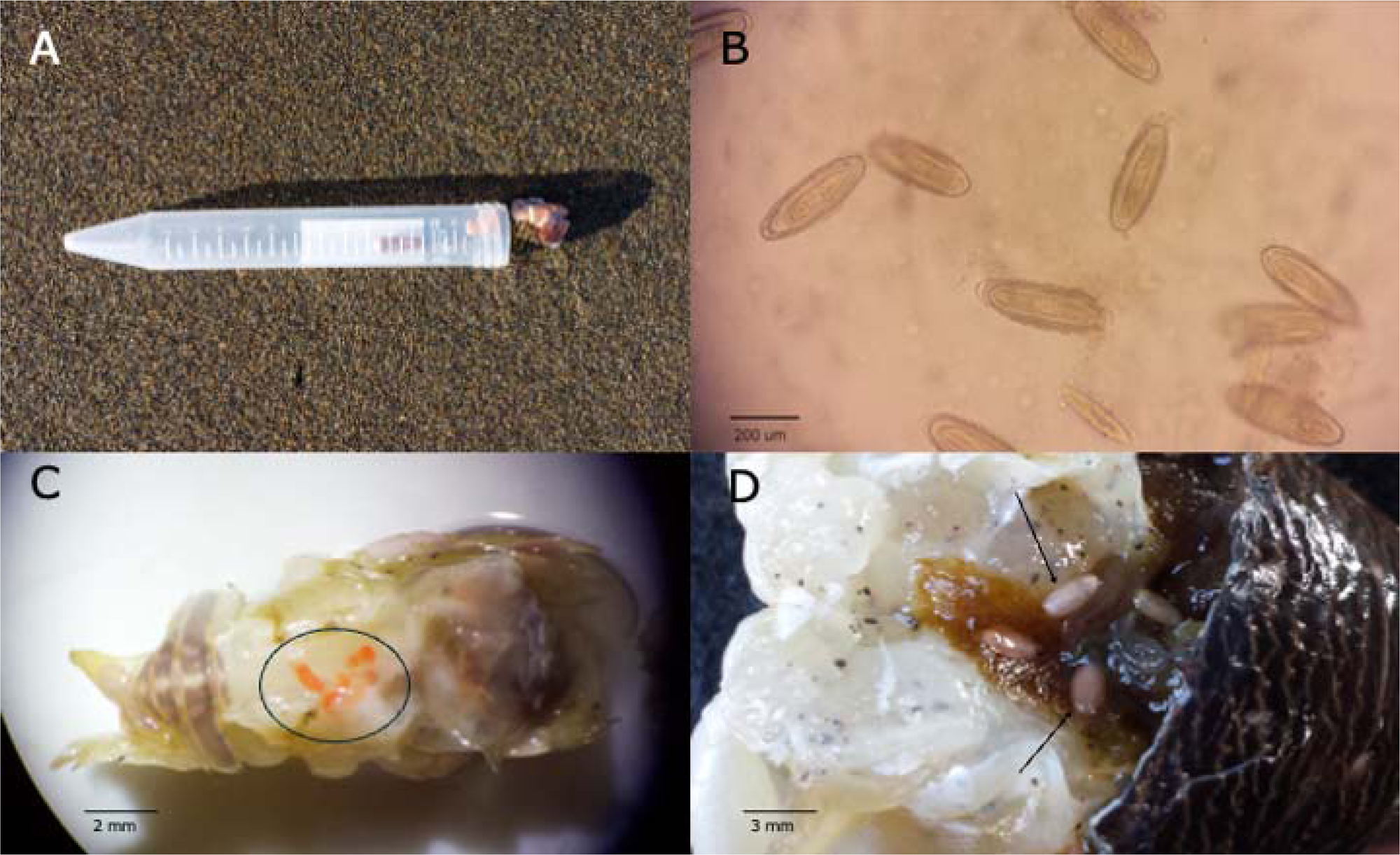
A = Faecal sample from grey gull *Leucophaeus modestus* in Curiñanco beach, Chile. B = Eggs of *P. altmani* extracted from faeces of the grey gull. Scale bar: 200 μm. C = Acanthellae of *P. altmani* in the hemocoel of the mole crabs *E. analoga* (cephalotorax removed). Circle indicates enclosed acanthellae. Scale bar: 2 mm. D = Cystacanths of *P. altmani* in the hemocoel of the mole crab, *E. analoga* (Cephalotorax removed). Rows indicate cystacanths. Scale bar: 3 mm. (Photography in Smith & Rodríguez, 2025).

**Figure 2:**
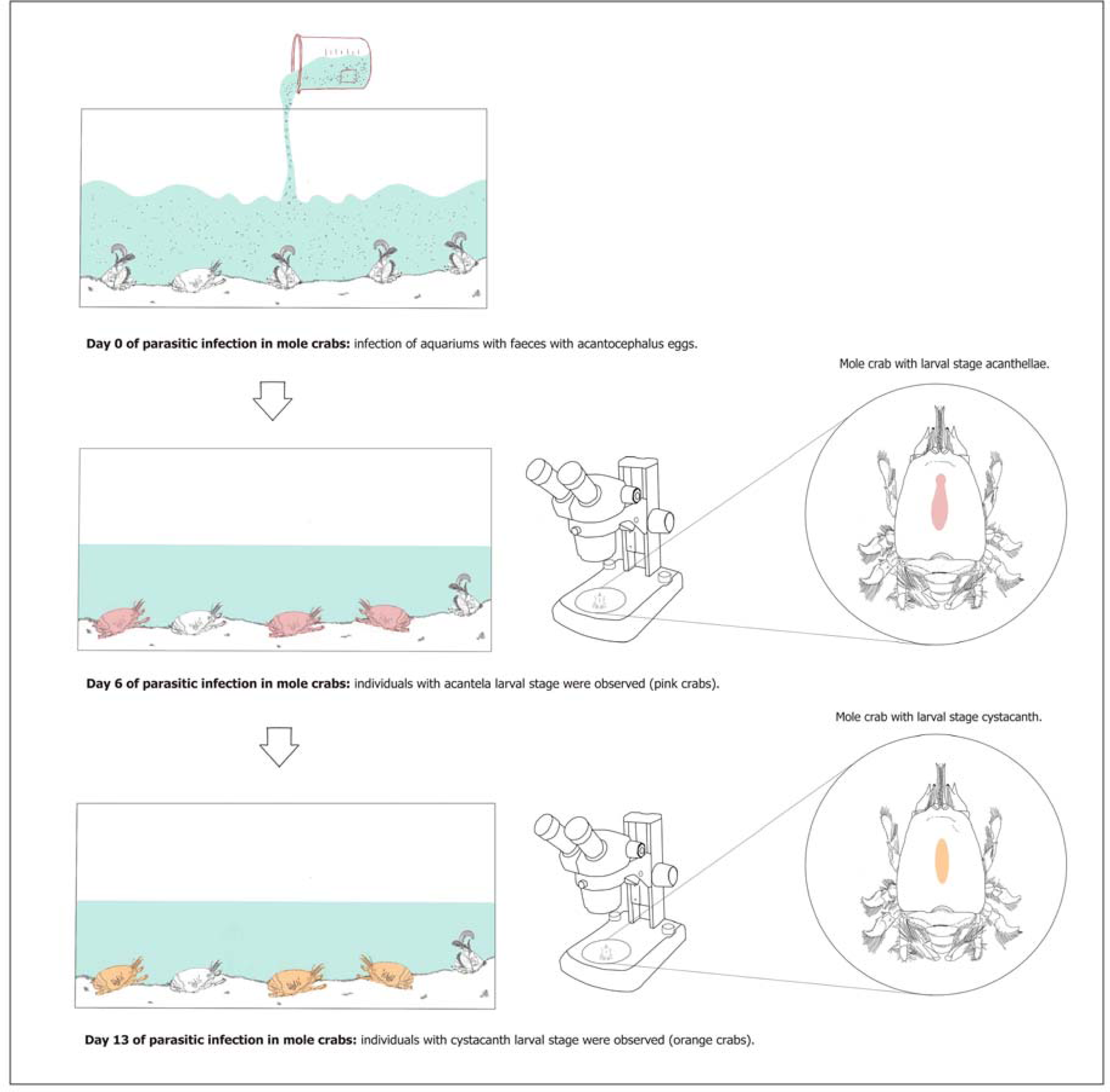
Experimental methodology of the egg’s inoculation. A = Inoculation of acanthocephalan *Profilicollis altmani* eggs obtained from seagull’s faeces. B = Detection of the acanthella larvae stage in the crustacean *Emerita analoga* from 7 days after inoculation. C = Detection of the cystacanth larval stage in the crustacean *Emerita analoga* from 13 days after inoculation.

The eggs of *P. altmani* in the faeces were detected using a microscope (4x; Presswell & Lagrue, 2016). The eggs appeared orange in color, with an elongated oval shape, typically resembling acanthocephalan eggs as described in previous studies (Smith & Rodríguez, 2025). From each faeces, three little portions were extracted, which were diluted in 20 ml marine sterilized water each one. Each portion was analysed to corroborate the presence, and measure the abundance of, parasite’s eggs (Fig. 1B).

### 2.3 Infection procedure

Prior to infection, all mole crabs were deprived of food for 24 h. Aquaria designated for infection were inoculated with 1.8 g of faeces diluted in 0.3 L of filtered seawater. Control aquaria were inoculated with the same amount of cleaned sand and filtered seawater. After 24 h, the water was replaced in all aquaria. Six days after inoculation, thirty mole crabs from each aquarium were dissected and inspected under a binocular microscope to detect and count acanthocephalan larvae (acanthellae and cystacanths) (Fig. 1C-D). Dissections were repeated on day thirteen. On day twenty-one, a second inoculation was performed with the same quantity of faeces (1.8 g in 0.3 L), following the initial protocol. Thirty days after the first inoculation, 30 mole crabs from each aquarium were dissected and inspected again, with a final dissection on day thirty-four. During each dissection, the cephalothorax length (mm) of each mole crab was measured to account for growth during the acclimatization period. The presence and number of acanthellae and cystacanths per host were recorded (Fig. 1C-D). Parasitological descriptors were calculated following Bush et al. (1997) and Rózsa et al. (2000): Prevalence was defined as the proportion of hosts infected with at least one parasite; mean abundance as the average number of parasites per host, including both infected and uninfected individuals; and mean intensity as the average number of parasites per infected host. Moreover, we used cystacanth occurrence (either present or absence) and abundance (number of individuals per host) in the formal statistical analyses.

### 2.4 Statistical analyses

We used information theory to assess the effects of parasite inoculation, time, host’s body size, and their interactions on parasitosis (Burnham, 2002). The candidate models included both separate and interactive effects on parasitosis (Table 1). Cystacanth occurrence and abundance were analysed with binomial (logit link) and Poisson (log link) generalised linear models, respectively. Model parameters were estimated through maximum likelihood. Bias-corrected Akaike Information Criteria (AICc) was used to select the model with the highest empirical support. Model selection was based on Δ*_i_*, expressed a AICc_i_ - AICc_min_, Akaike weight, which represents the probability of each model, model as 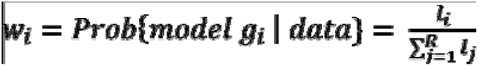, where 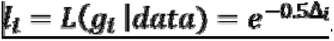. The evidence ratio of the best-supported model was calculated as *w_top_* / *w_j_* (Burnham, 2011). A likelihood ratio based pseudo-coefficient of determination (R^2^) was calculated for each top model. Since acanthellae are still in early development and unable to infect the definitive host, they were excluded from the statistical analysis (Balboa Figueroa et al., 2019).

**Table 1.**
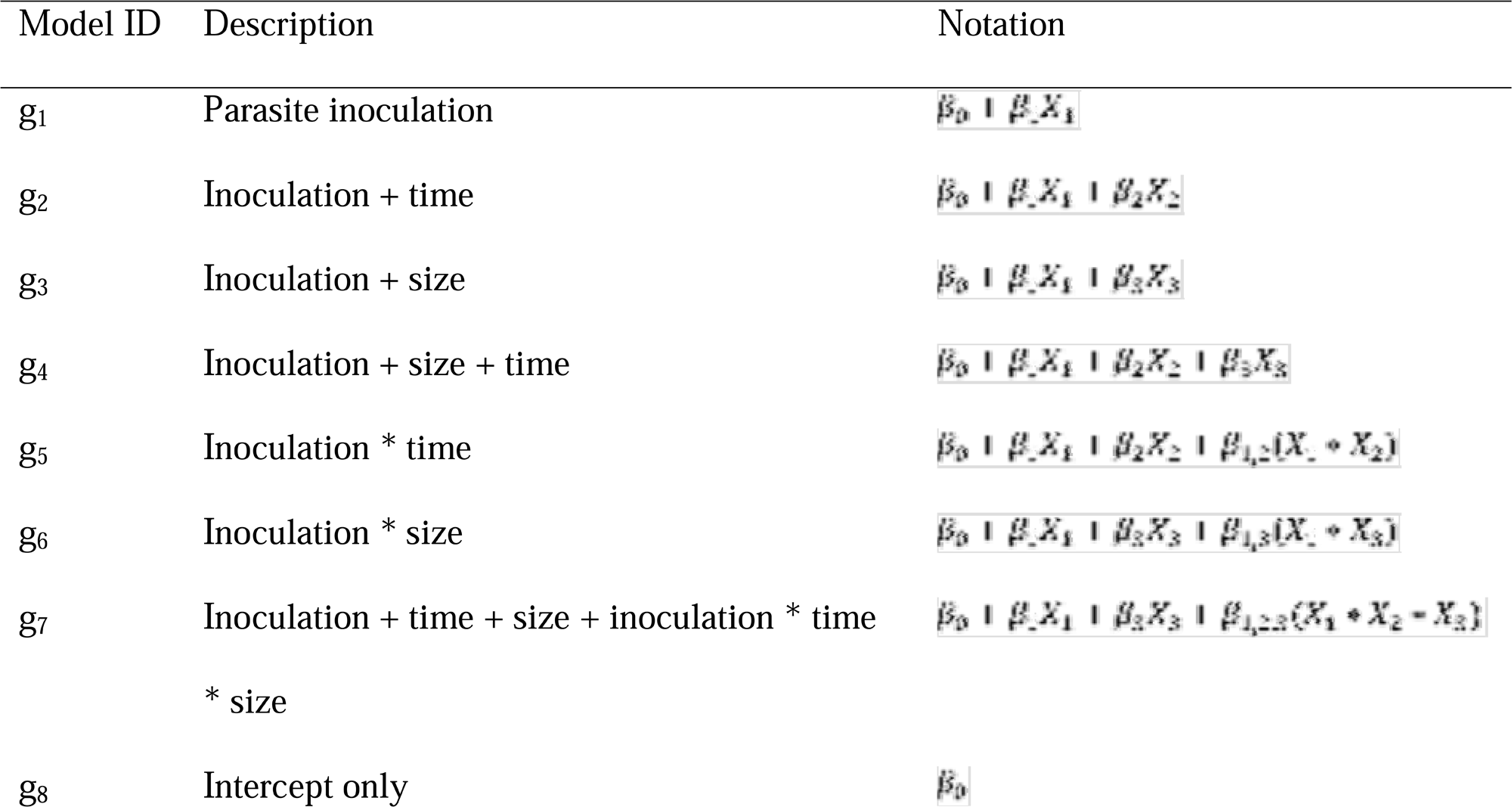
Model set used to evaluate the effects of parasite inoculation, time, and host body size on infection status and intensity. The models vary in complexity, including additive and interaction terms among inoculation treatment, time post-inoculation, and host size. The notation column indicates the explanatory variables included in each model: “+” indicates additive effects, and “*” indicates interactions.

All statistical analyses were undertaken in Program R 4.0.1 (R Development Core Team, 2020). Linear mixed-effects models were computed with the *lme4* (Bates et al., 2015).

## RESULTS

### Larval development: transition from acanthellae to cystacanths across two inoculation events

Before the first inoculation, no mole crab was infected with acanthellae and only few (i.e., five) were infected with cystacanths (Fig. 3A and B). After the first inoculation, the prevalence of acanthellae in large mole crabs was generally higher than in small crabs (Fig. 3A). In contrast, mean abundance was similar between both body-size groups (Fig. 4A and 4B). The acanthella stage emerged on the sixth day after inoculation of parasite eggs (Fig. 3A, 4A). Six days after inoculation, 83.3% of the small mole crabs were infected (Fig. 3A and 3B). Mean abundance and mean intensity were 1.2 and 1.4 acanthellae *per* host at day six (Fig. 4A). On day six, the large mole crabs showed 96.6% of prevalence, and 1.4 and 1.5 of mean abundance and intensity, respectively (Figs. 3 and 4).

**Figure 3:**
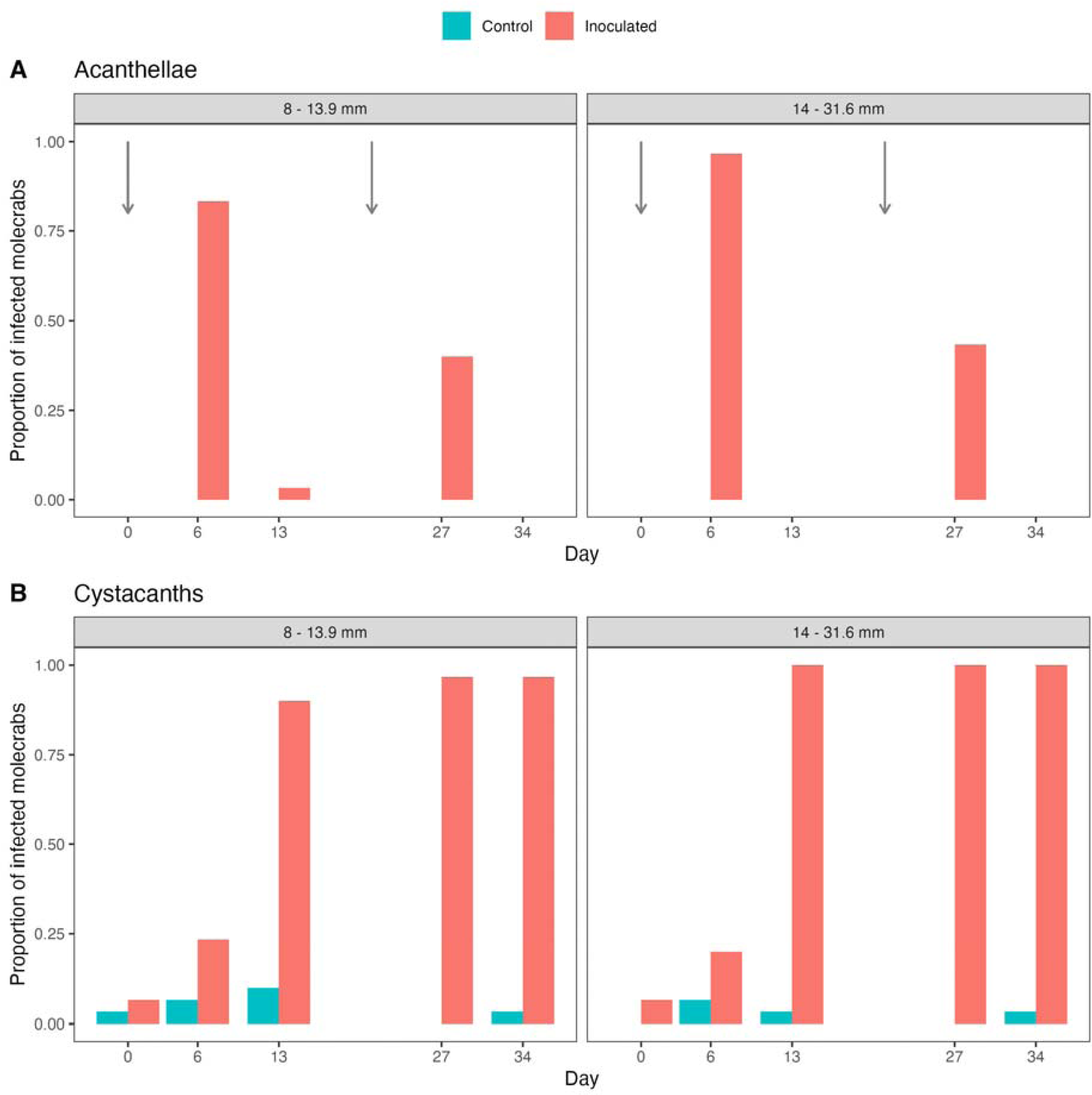
Proportion of infected molecrabs by acanthocephalan parasites during the egg inoculation experiment in two host body sizes groups over the course of the experiment. A = Proportion of infected molecrabs by acanthellae larvae following egg inoculation. B = Proportion of infected molecrabs by cystacanths larvae following egg inoculation. The black arrow indicates the day on which parasite egg inoculation was performed. The first inoculation took place at “Day 0” and the second at day 21.

**Figure 4:**
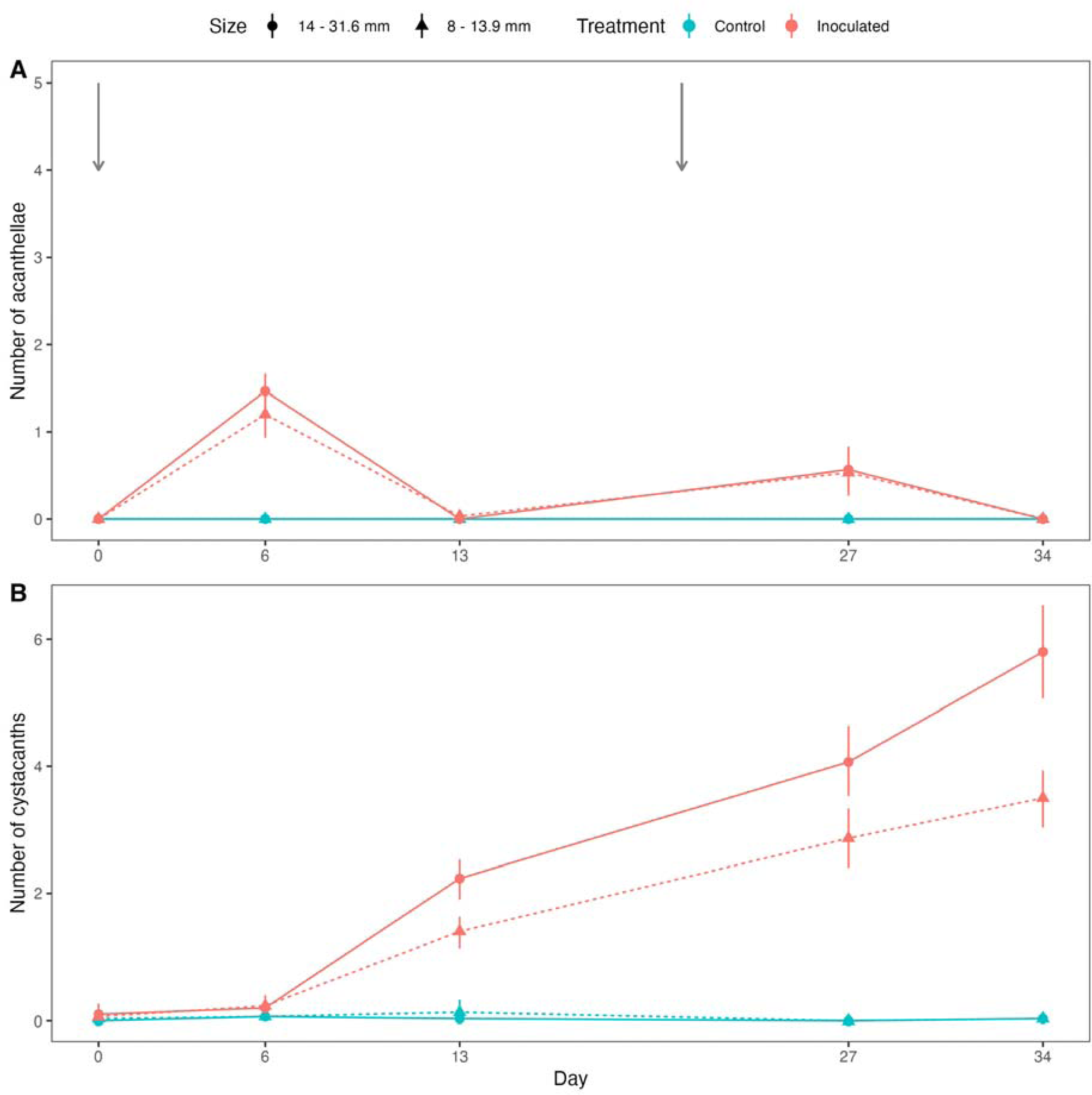
Number of acanthocephalan parasites (abundance) during the egg inoculation experiment in two host body sizes groups over the course of the experiment. A = Number of acanthellae larvae developed in *Emerita analoga* following egg inoculation. B = Number of cystacanths larvae developed in *Emerita analoga* following egg inoculation. The black arrow indicates the day on which parasite egg inoculation was performed. The first inoculation took place at “Day 0” and the second at day 21.

These acanthellae rapidly matured into the cystacanth stage (Fig. 3B). By day 13, most acanthellae had disappeared, coinciding with a marked increase in cystacanth prevalence (Fig. 3B). Small crabs exhibited 90% prevalence, with mean abundance and intensity of 1.4 and 1.5 cystacanths *per* host, respectively; large crabs reached 100% prevalence with mean abundance and intensity of 2.2 cystacanths per host each, confirming successful parasite development within the hosts (Fig. 3B and 4B).

The second inoculation, taking place on day 21, resulted in an increased parasitosis across crab’s body sizes (Fig. 4). For example, by day 27, small crabs showed 40% prevalence of acanthellae, with mean abundance and intensity of 0.5 and 1.3 acanthellae per host, respectively (Figs. 3 and 4). Large crabs exhibited 43.4% prevalence, with mean abundance and intensity of 0.56 and 1.3 acanthellae per host, respectively (Figs. 3 and 4). As in the first infection cycle, these acanthellae matured rapidly: by day 34, acanthellae were no longer observed, and cystacanths predominated (Fig. 3 and 4). At this moment, small crabs reached 97.6% prevalence of cystacanths, with mean abundance and intensity of 3.5 and 3.6, cystacanths per host respectively; large crabs reached 100% prevalence, with mean abundance and intensity both at 5.8 cystacanths per host (Fig. 3 and 4).

The analysis of model selection supported the effectiveness of the experimental inoculation of parasite eggs. For the probability of occurrence of cystacanths (for brevity, referred to as infection probability hereafter), the model with the highest empirical support (lowest AICc) was that including the separate and interactive effects of inoculation, time, and host’s body size (*g_7_* in Table 1). The probability of *g_7_* of being the best model given the data (Akaike weight, *w_7_*) and model set was 0.97, while that of the model with the closest Δ*_i_*was 0.03 (*g_5_*, Δ*_5_* = 6.94; Table 2). Thus, the empirical support for model *g_7_* was ca. 32 times that of the closest competing model (evidence ratio = 32.1). Importantly, *g*_7_ was 2.9 * 10^112^ more likely than the null model (*g_8_* in Table 2), which provides a very strong empirical support to the time-, and body size-dependent effect of the inoculation on the probability of infection. Model *g*_7_ accounted for an 81 % of the variation in infection probability (pseudo-*R^2^* = 0.81). The predicted probability of infection steeply increased over time as a result of the inoculation; this increment was faster as host’s body length increased (Fig. 5A, Table 4).

**Figure 5:**
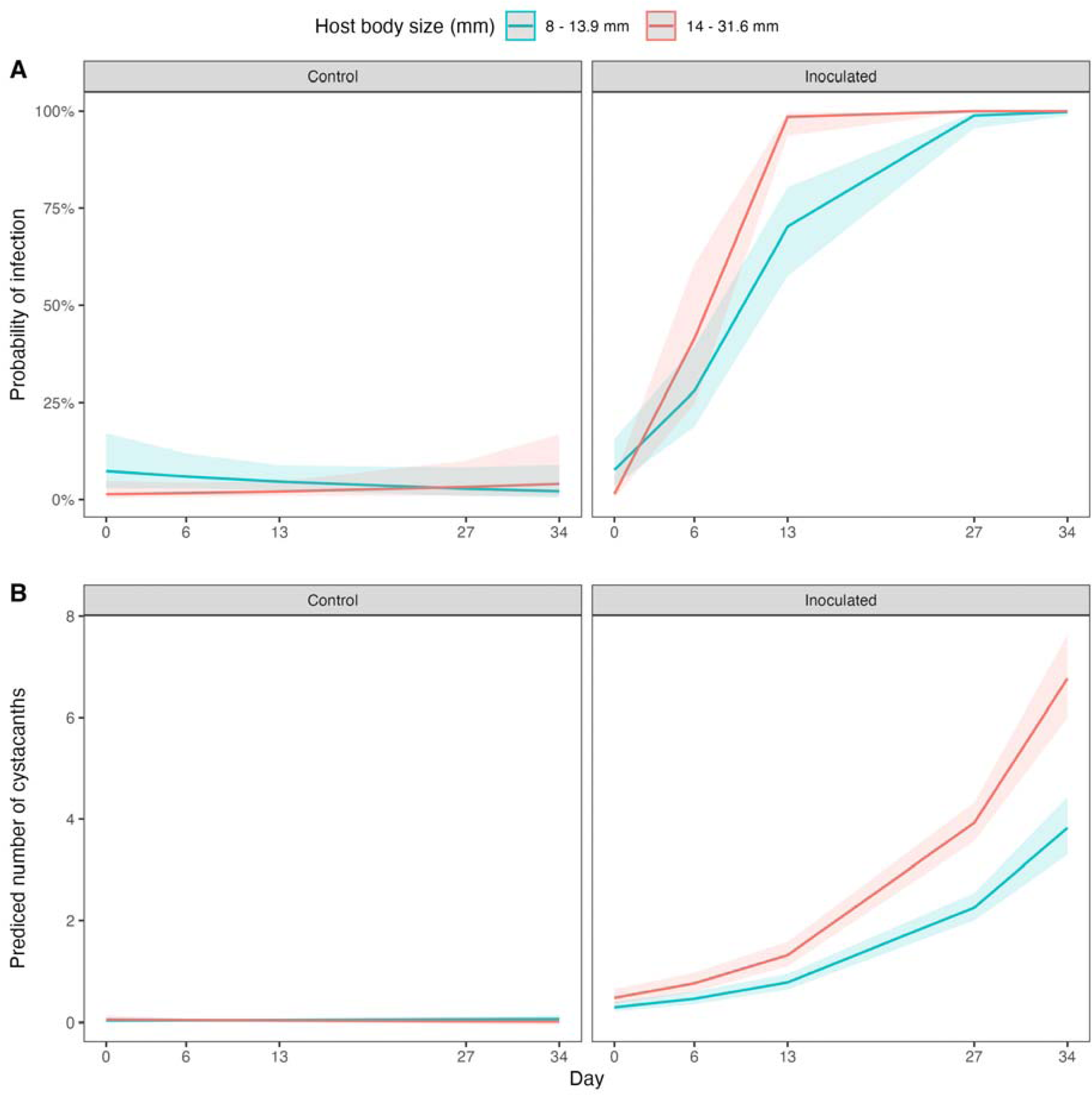
A = Probability of cystacanth infection in molecrabs as a function of host body size, before and after parasite egg inoculation, over time. B = Predicted number of cystacanths in molecrabs as a function of host body size, before and after parasite egg inoculation, over time

**Table 2.** Summary of information-criteria-based model selection for the probability of infection (occurrence of cystacanths) in mole crabs. Models include effects of inoculation (I), time (T), host body size (S), and their interactions. Columns show degrees of freedom (df), corrected Akaike Information Criterion (AICc), difference in AICc relative to the best model (Δ), Akaike weights (w), and evidence ratios (ER). The best-supported model (g7) includes the three-way interaction (I:T:S), indicating that inoculation effects on infection probability vary with time and host size.

| Mod | I | T | S | I:T | I:S | I:T:S | df | AICc | $\Delta$ | $l$ | $w$ | ER |
| --- | --- | --- | --- | --- | --- | --- | --- | --- | --- | --- | --- | --- |
| $g_7$ | + | + | + | | | + | 6 | 252.00 | 0.00 | $1.00 * 10^0$ | $9.70 * 10^{-1}$ | $1.00 * 10^0$ |
| $g_5$ | + | + | | + | | | 4 | 258.94 | 6.94 | $3.11 * 10^{-2}$ | $3.02 * 10^{-2}$ | $3.21 * 10^1$ |
| $g_2$ | + | + | | | | | 3 | 329.07 | 77.07 | $1.84 * 10^{-17}$ | $1.79 * 10^{-17}$ | $5.45 * 10^{16}$ |
| $g_4$ | + | + | + | | | | 4 | 329.95 | 77.94 | $1.19 * 10^{-17}$ | $1.15 * 10^{-17}$ | $8.42 * 10^{16}$ |
| $g_3$ | + | | + | | | | 3 | 490.29 | 238.29 | $1.81 * 10^{-52}$ | $1.75 * 10^{-52}$ | $5.54 * 10^{51}$ |
| $g_1$ | + | | | | | | 2 | 490.39 | 238.39 | $1.72 * 10^{-52}$ | $1.66 * 10^{-52}$ | $5.83 * 10^{51}$ |
| $g_6$ | + | | + | | + | | 4 | 491.44 | 239.44 | $1.02 * 10^{-52}$ | $9.85 * 10^{-53}$ | $9.84 * 10^{51}$ |
| $g_8$ | | | | | | | 1 | 769.92 | 517.91 | $3.44 * 10^{-113}$ | $3.34 * 10^{-113}$ | $2.91 * 10^{112}$ |

The number of cystacanths per host—i.e., abundance—systematically responded to the experimental inoculation over time. The model including the separate and interactive effects of inoculation, time, and host’s body size scored the best empirical support, evidenced by the lowest AICc (*g_7_* in Table 3). The Akaike weight of *g_7_*was 0.99, and this model was 166 times more likely than the closest competing model (*g_4_,* Δ*_4_* = 10.22, *w_4_* = 0.006, evidence ratio of *g_7_* vs. *g_4_* = 165.7). The empirical evidence for *g7* was 3.34 * 10^276^ times that of the null model (*g_8_*in Table 3), providing a very conclusive support to the effect of the inoculation on intensity. Model *g*_7_ accounted for an 91 % of the variation in infection probability (pseudo-*R^2^* = 0.91). The predicted intensity steeply and nonlinearly increased over time as a result of the inoculation and this increment was faster for larger hosts (Fig. 5B, Table 4).

**Table 3.** Summary of information-criteria-based model selection for parasite intensity (number of cystacanths per host) in mole crabs. Models include the effects of inoculation (I), time (T), host body size (S), and their interactions. Table shows degrees of freedom (df), corrected Akaike Information Criterion (AICc), difference in AICc relative to the best model (Δ), likelihood (l), Akaike weights (w), and evidence ratios (ER). The best-supported model (g7) includes the three-way interaction (I:T:S), indicating that the intensity of infection depends on the combined effects of inoculation, time, and host size.

| Mod | I | T | S | I:T | I:S | I:T:S | df | AICc | $\Delta$ | *l* | *w* | *ER* |
| --- | --- | --- | --- | --- | --- | --- | --- | --- | --- | --- | --- | --- |
| g7 | + | + | + | | | + | 6 | 899.36 | 0 | $0.99 * 10^0$ | $1.00 * 10^0$ | $1.00 * 10^0$ |
| g4 | + | + | + | | | | 4 | 909.58 | 10.22 | $6.00 * 10^{-3}$ | $6.00 * 10^{-3}$ | $1.65 * 10^{-2}$ |
| g5 | + | + | | + | | | 4 | 944.53 | 45.17 | $1.55 * 10^{-10}$ | $1.56 * 10^{-10}$ | $6.43 * 10^9$ |
| g2 | + | + | | | | | 3 | 959.41 | 60.05 | $9.06 * 10^{-14}$ | $9.11 * 10^{-14}$ | $1.1 * 10^{13}$ |
| g6 | + | | + | | + | | 4 | 1358.10 | 458.74 | $2.4 * 10^{-100}$ | $2.4 * 10^{-100}$ | $4.1 * 10^{99}$ |
| g3 | + | | + | | | | 3 | 1358.64 | 459.28 | $1.8 * 10^{-100}$ | $1.9 * 10^{-100}$ | $5.4 * 10^{99}$ |
| g1 | + | | | | | | 2 | 1425.67 | 526.31 | $5.1 * 10^{-115}$ | $5.2 * 10^{-115}$ | $1.9 * 10^{114}$ |
| g8 | | | | | | | 1 | 2172.80 | 1273.44 | $3 * 10^{-277}$ | $3 * 10^{-277}$ | $3.3 * 10^{276}$ |

**Table 4.** Summary of best (K-L) models for infection probability and parasite intensity (cystacanths). Estimates, standard errors (SE), and z-values for fixed effects are shown, including inoculation treatment, time, host body size, and their interactions. These models explain how these factors influence the presence and number of cystacanth parasites in mole crabs.

|  | Infection probability |  |  | Intensity |  |  |
| --- | --- | --- | --- | --- | --- | --- |
|  |  |  | z- |  |  | z- |
|  | Estimate | SE | value | Estimate | SE | value |
| Intercept | -0.93 | 0.94 | -0.99 | -3.86 | 0.54 | -7.15 |
| Inoculation | 0.04 | 0.61 | 0.07 | 2.18 | 0.45 | 4.80 |
| Time | -0.10 | 0.06 | -1.58 | 0.07 | 0.01 | 5.99 |
| Size | -0.15 | 0.06 | -2.56 | 0.04 | 0.02 | 2.20 |
| Control:Time:Size | 0.01 | 0.00 | 1.57 | 0.00 | 0.00 | -2.92 |
| Inoculated:Time:Size | 0.03 | 0.01 | 5.10 | 0.00 | 0.00 | 0.30 |

## DISCUSSION

This study demonstrated that the experimental inoculation with fresh gull faeces carrying eggs of the acanthocephalan *Profilicollis altmani* enables effective, predictable, and repeatable infections in marine hosts maintained under laboratory conditions. We successfully infected mole crabs (*Emerita analoga*) of different sizes, observing larval stages from day six onwards and a progressive accumulation of cystacanths up to day thirty-four. The statistical models that best explained infection probability and abundance, therefore, included the interactive effects of treatment, time, and host body size, showing high explanatory power (pseudo-R² > 0.80). These results validate this methodology as a strong experimental alternative for studying parasites with complex life cycles, while also allowing manipulation of key variables that are difficult to control in field conditions. This technique, based on administering parasite eggs naturally present in the faeces of the definitive host, emerges as a robust tool for numerically balancing experimental groups, allowing for the availability of both infected and uninfected hosts in controlled proportions. Based on these findings, we will discuss (i) the relevance of using controlled infection models in parasitic ecology, (ii) the limitations of natural sampling in systems with cryptic or heterogeneous infections, (iii) the ecological value of statistically homogenizing parasite burden, and (iv) the potential of this approach for future experimentation in ecophysiology, behavior, and ecotoxicology.

One of the major challenges in studying parasites with complex life cycles lies in obtaining naturally infected and uninfected hosts across their entire ontogenetic range, as larger individuals tend to accumulate higher parasite loads (Ebert, 1999; Rodríguez & Valdivia, 2017; Lorenti et al., 2018; Rodríguez et al., 2022). In this experiment, only presumptively uninfected juvenile hosts (<9 mm; Rodríguez & Valdivia, 2017) were collected and reared under controlled conditions until they reached larger sizes, with the aim of eliminating this source of bias. Nevertheless, it is important to note that some crabs were naturally infected prior to collection, despite being selected according to the previously reported minimum infection size threshold (≥9 mm cephalothorax length) (Rodríguez & Valdivia, 2017). This experimental approach allowed us to isolate the effect of body size on infection, revealing that larger hosts not only acquired more intense infections but did so in a shorter period of time. This pattern can be explained by functional mechanisms associated with feeding. *Emerita analoga* is an active filter feeder, and larger individuals process significantly greater volumes of water and sediment than smaller conspecifics (Dugan & Hubbard, 1996; Lastra et al., 2004). Consequently, higher filtration activity increases the encounter rate between the host and parasite eggs present in the water column, thereby enhancing the probability of ingesting infective stages. In contrast, smaller mole crabs filter reduced volumes of water and are therefore exposed to lower effective densities of parasite eggs, which could account for the lower parasite burdens observed following inoculation (Smith, 2007; Balboa et al., 2009; Rodríguez & Valdivia, 2017). This trend was confirmed in our experiment, where both size classes were inoculated simultaneously, allowing us to disentangle the effects of host age from those associated with filtration rate on parasitism. Overall, these findings indicate that body size, beyond representing ontogenetic age, acts as a mechanistic determinant of parasite exposure, opening new perspectives for integrating morpho-functional variables into transmission models in natural systems.

The main novelty of this study lies in the use of fresh faeces from definitive hosts as a vector to induce infections in intermediate hosts. While faeces are routinely employed for parasite diagnosis (Mas-Coma et al., 2014; Presswell & Lagrue, 2016; Rodríguez & George-Nascimento, 2021), their application as an experimental tool in ecological studies is almost non-existent. This technique leverages the natural faecal-oral transmission route of the parasite and circumvents the complex procedures involved in isolating eggs or larvae (Dianne et al., 2012; Presswell & Lagrue, 2016). By enabling a realistic and efficient inoculation without the need for direct manipulation of the parasites, its use is facilitated especially in marine organisms with opaque cuticles where visual detection of infection is impossible (Balboa Figueroa et al., 2019). For example, the transparent cuticula of the amphipod *Gammarus pulex* allows one to visually detect the presence of acanthocephalan cystacanths without dissection (Bakker et al., 1997; Thünken et al., 2010, 2018). In contrast, *E. analoga* possesses an opaque cuticle that prevents any external visual assessment of infection status, rendering non-invasive approaches unfeasible. In this context, the use of faeces as an experimental vector represents an effective, standardized, and ecologically relevant alternative for inducing infections in systems with technical diagnostic limitations. Therefore, this methodology expands the possibilities to study parasitic systems where infection is cryptic and difficult to standardize.

The methodology presented here not only represents a technical innovation but also opens new avenues for the development of controlled experiments in disease ecology. In systems involving parasites with complex life cycles, where infection occurs in a cryptic and (seemly) stochastic manner, controlling the infection status of hosts has remained a persistent challenge (Balboa et al., 2009; Rodríguez et al., 2017; Rodríguez et al., 2022). Our technique allows for the standardization of initial experimental conditions by generating controlled groups of infected and uninfected individuals, ensuring treatment replicability and a balanced distribution of samples across experimental groups (Albert et al., 2010; Burgess et al., 2022). This is particularly valuable in studies aiming to assess the effects of parasites on host phenotypic traits such as behavior, metabolic rate, growth, or survival, where natural infection biases could confound result interpretation (Franceschi et al., 2008; Thünken et al., 2010; Goedknegt et al., 2019). Moreover, this technique is scalable to other marine or aquaculture systems where experimental evaluation of parasite impacts under semi-controlled or laboratory conditions is desired. Its application can extend to studies on co-infections, parasite competition, changes in host life-history traits, or parasite-mediated trophic interactions, contexts in which sample size balancing is often difficult due to the nature of individual life cycles (Balboa et al., 2009; Goedknegt et al., 2019 Thünken et al., 2019). Additionally, this approach is not limited to acanthocephalans but could be applied to other parasite groups with complex life cycles whose infective stages are also released through the faeces of definitive hosts. These include nematodes, cestodes, and digeneans, where eggs or larvae are expelled by aquatic or semi-aquatic vertebrates and subsequently ingested by intermediate hosts. Therefore, experimental inoculation via faeces represents an ecological, efficient, and replicable pathway to induce infections across a wide diversity of parasitic systems.

Despite its advantages, the faecal inoculation method presented here has inherent limitations that should be acknowledged. First, the use of fresh faeces as the source of parasite eggs introduces variability in the exact number and viability of eggs ingested by each host, potentially leading to uncontrolled variation in infection doses among individuals (Dianne et al., 2011). This contrasts with methods using isolated and quantified parasite stages, which allow for precise dosing. Additionally, faecal suspensions may contain a mixture of parasite species and developmental stages, which could confound interpretations if the parasite community is diverse (Mas-Coma et al., 2014). Moreover, environmental factors affecting egg viability and infectivity within the faecal material, such as temperature or microbial activity, could influence infection success and are challenging to standardize fully (Presswell & Lagrue, 2016; Hopkins et al., 2022). Lastly, although the method mimics natural transmission routes, it does not fully replicate the complexity of environmental exposure in situ, such as spatial heterogeneity of eggs, host behavior, and ecological interactions that may influence infection dynamics in the wild (Ebert 1999; Balboa et al., 2009). Therefore, while this technique provides a robust framework for controlled experimental infections, careful consideration of these limitations is essential when extrapolating laboratory findings to natural systems.

## CONCLUSION

This study proposes and validates a simple, effective, and ecologically realistic experimental tool for inducing parasitic infections in intermediate hosts through the inoculation of fresh faeces containing viable parasite eggs. Using *Emerita analoga* as the model host and the acanthocephalan *Profilicollis altmani* as the parasite, we demonstrate that it is possible to overcome common methodological limitations in ecological studies of parasites with complex life cycles, particularly the difficulty of obtaining naturally infected and uninfected individuals of different sizes. Our results show that, under controlled conditions, larger hosts acquired higher infection intensities, likely due to their greater filtration capacity and cumulative larval exposure over time. This technique, being consistent with the parasite’s natural faecal-oral transmission route and avoiding the need for direct manipulation of infectious propagules, offers substantial advantages in terms of replicability, experimental control, and applicability to systems with cryptic or hard-to-detect infections. Furthermore, its use may extend beyond acanthocephalans to other parasitic groups with similar life cycles, such as nematodes, cestodes, and digeneans. Taken together, we propose that this methodology represents a significant experimental advance for the design of ecological studies, enabling the formulation of *a priori* hypotheses, infection standardization, and improved statistical inference, critical components for understanding the influence of parasites on host life-history traits, behavior, and population dynamics.

## ACKNOWLEDGEMENTS

We say thanks to BEL Laboratory at Universidad Austral de Chile for supporting in the field and experimental work.

## ADDITIONAL INFORMATION AND DECLARATIONS

### Author contributions

SMR and NV conceived the ideas, designed methodology, and analysed the data. SMR KBA, BG, and VE-A conducted the research. SMR led the writing of the manuscript. All authors contributed critically to the drafts and gave final approval for publication.

### Data availability

All data generated by the postdoctoral research is open access and available via Dryad Digital Repository XXX

---

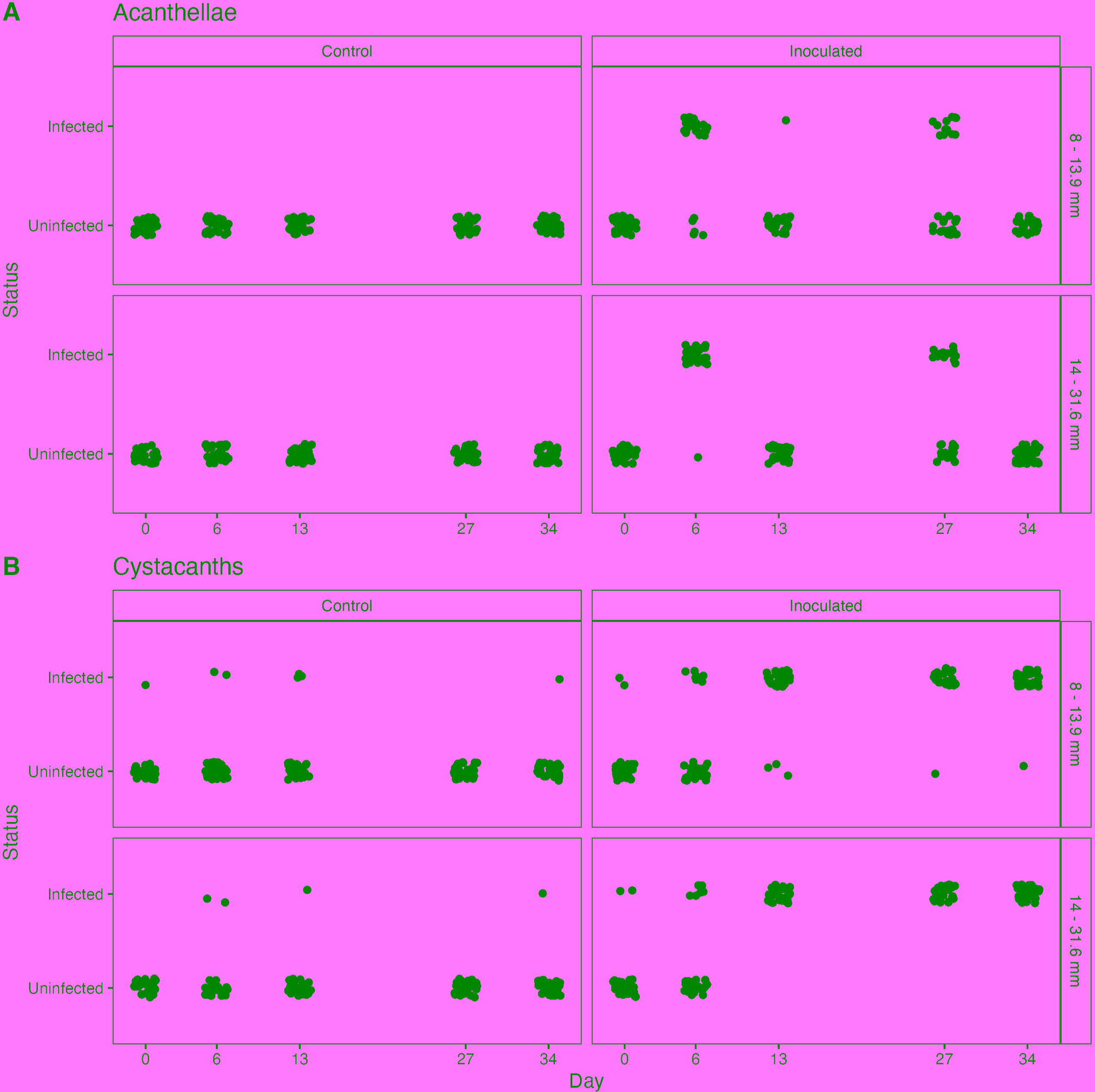

